# Septal GABAergic Inputs to CA1 Govern Contextual Memory Retrieval

**DOI:** 10.1101/824748

**Authors:** Arnau Sans-Dublanc, Adrià Razzauti, Srinidhi Desikan, Marta Pascual, Jaime de la Rocha, Hannah Monyer, Carlos Sindreu

## Abstract

Whether projections from the medial septum regulate the function of the CA1 hippocampus in episodic memory retrieval is not known. Here we show that septal GABAergic inputs to CA1 promote contextual fear memory, blocking the activation of parvalbumin-rich interneurons to facilitate Erk/MAP-kinase signaling in pyramidal cells during retrieval. Thus, suppression of feed-forward inhibition onto CA1 by septal GABAergic neurons gates contextual fear behavior.

## Introduction

Memory retrieval, or the access to stored information in the brain, guides several adaptive behaviors^1^, and its precision or success is likely to be impaired during dementia or autism^2,3^. Although the distributed networks supporting retrieval are not well understood, studies point to the CA1 output region of the hippocampus as a critical node for the recall of episodic memory, such as contextual conditioning^4–6^. In this paradigm, retrieval is reliably inferred from the rapid freezing behavior upon re-exposure, triggering context-specific signatures at the cell firing^7^ or molecular levels^8^. Considerable progress has linked CA1 pyramidal cell activation with afferences in CA3 or entorhinal cortex^9–11^, and with downstream circuits in the subiculum^12^ or neocortex^13^ during contextual recall. However, the hippocampus also receives long-range GABAergic projections from the subcortical medial septum / diagonal band (i.e. MS) that terminate on interneurons^14^, convey sensory stimuli^15^, and degrade with aging^16^. MS^GABA^ inputs disinhibit hippocampal pyramidal cells^17^, and contribute to rhythmic oscillations necessary during REM sleep for memory consolidation^18^. Interestingly, computational models predicted that septal GABAergic inhibition can facilitate memory retrieval by occluding feed-forward interneurons^19^, but this hypothesis has not been investigated.

## Results

To examine the function of the MS–CA1 pathway, we first injected into the dorsal CA1 region a herpes simplex virus expressing YFP (HSV-YFP) for high efficiency brain-wide retrograde cell tagging (Supplementary Fig. 1a-c). Up to 7 ± 1 % of CA1-projecting cells originated from the MS (Fig. 1a). MS^CA1^ cells were predominantly GABAergic (~86%) and, in dorsal and medial divisions, they concentrated within ~ 200 μm from midline, where the ratio of GABAergic-to-cholinergic cells was highest (Supplementary Fig. 1d,e). Injecting the CA1 with a retrograde virus expressing Cre recombinase, CAV2-cre, followed by injection into the MS of AAV-Flex-tdTomato for conditional expression of red fluorescent protein selectively labeled MS–CA1 projections (Fig. 1b). Axons of MS^CA1^ cells mainly targeted the CA1, dorsal subiculum or lateral perforant path, indicating little collateralization (Supplementary Fig. 2a,b). Consistently, co-labeling of MS cells projecting to the entorhinal cortex, another known target, revealed intermingled but segregated subpopulations (Supplementary Fig. 2c,d). To test whether the MS–CA1 pathway is recruited during memory retrieval, we trained mice for contextual fear conditioning and quantified the presence of c-Fos, a marker of recent activation, in retrogradely tagged cells. Contextual re-exposure to evoke retrieval produced marked freezing behavior (Supplementary Fig. 3a,b), and induced a significant increase in c-Fos expression compared with controls (Fig. 1c-d).

**Figure 1.**
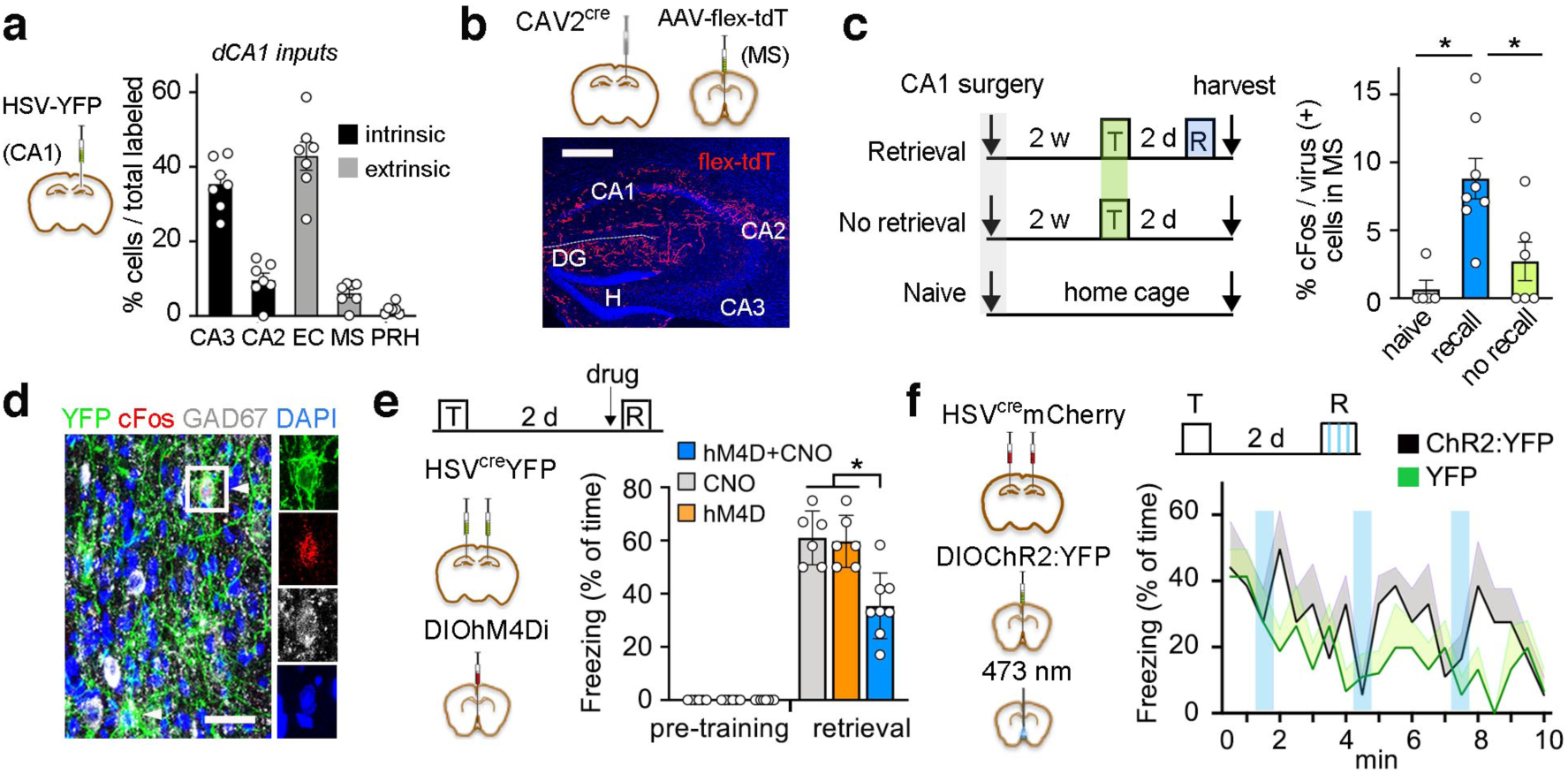
Projections from MS to CA1 promote retrieval of contextual fear memory. (**a**) Delivery of HSV-YFP into the CA1 labels afferent cell bodies (n = 7 mice). (**b**) Distribution of labeled axons originating from CA1-projecting septal (MS^CA1^) cells. Scale bar, 0.5 mm. (**c**) Experimental design and quantification of c-Fos in MS^CA1^ cells from naïve (n = 5), trained (T/no recall, n = 6) or retrieved (R/recall, n = 8) mice. Retrieval increased Fos immunoreactivity (F_(2,19)_ = 11.9, P = 0.001; post-hoc tests, P = 0.012 compared with naïve or P = 0.0007 with no recall). (**d**) Co-labeling of HSV-derived YFP, c-Fos and GAD-67 in MS after retrieval. Scale bar, 0.02 mm. (**e**) Chemogenetic inhibition of MS^CA1^ neurons (hM4D+CNO, n = 8) during contextual retrieval (effect of treatment F_(2,34)_ = 12.3, P < 0.0001; effect of conditioning F_(1,34)_ = 438, P < 0.0001; interaction F_(2,34)_ = 12.3, P < 0.0001; RM ANOVA followed by Sidak’s test, P < 0.0001 compared with controls, each n = 6). (**f**) Effect of photostimulation of MS^CA1^ expressing ChR2:YFP or YFP during re-exposure (n = 6 mice / group; effect of treatment F_(1, 12)_ = 11.15, P = 0.038; effect of time F_(1, 12)_ = 2.2, P = 0.045; interaction F_(1, 12)_ = 1.03, P = 0.42). Shaded colors indicate SD.

We expressed the inhibitor receptor hM4Di fused to mCherry in MS^CA1^ cells using a conditional approach as above (Supplementary Fig. 3c). Targeted neurons were then inhibited with injection of the agonist CNO before the retrieval test. Selective inhibition of MS^CA1^ neurons impaired fear memory compared with mice injected with vehicle or mice expressing YFP and receiving CNO (Fig. 1e). Conversely, MS^CA1^ neurons were transduced with channelrodhopsin (AAV-DIOChR2:EYFP) and their cell bodies photostimulated at 10 Hz (i.e. their natural frequency^18^) in 30 s epochs during contextual re-exposure. Although the fraction of c-Fos+ cells doubled (Supplementary Figs. 3d,g), light pulses only transiently increased freezing compared with YFP control, and both groups showed a gradual decline over the course of 10 minutes (Fig. 1f), indicating that the effect of physiological MS^CA1^ activation is near saturation.

The above results indicate that MS^CA1^ neurons are critical for memory recall, and suggest that GABAergic septal (MS^GABA^) terminals in CA1 can support such function. To directly test this hypothesis, we injected GAD2-cre mice with AAV-FlexPSAM^L141F^:GlyR to express the Cl-permeable, chimeric receptor exclusively in MS^GABA^ neurons, followed by bilateral cannulae in CA1 for local injection of its agonist PSEM (Fig. 2a). PSAM:Gly identified with α-bungarotoxin was highly expressed at distal septo-hippocampal terminals (Fig. 2b). PSEM infusion confined to CA1 (Supplementary Fig. 4a,b) impaired memory retrieval compared with controls (Fig. 2c), supporting the conclusion that the MS^GABA^–CA1 pathway is required for contextual recall.

**Figure 2.**
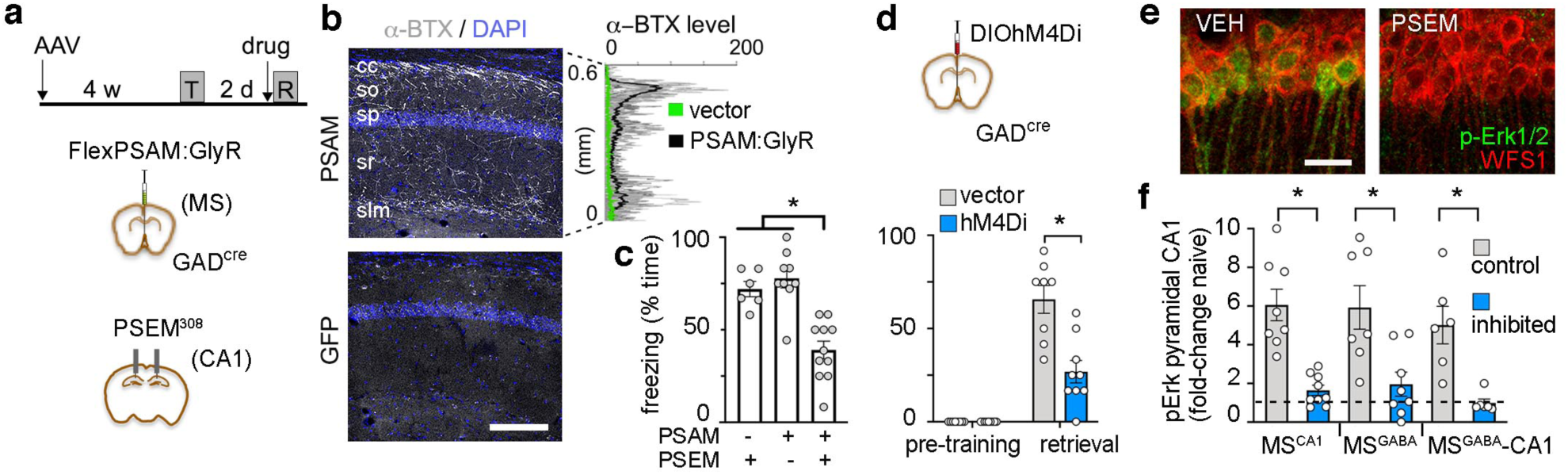
Septal GABAergic afferences in recall and associated Erk1/2 activation. (**a**) Experimental design. (**b**) Bungarotoxin fluorescence (α-BTX) showing PSAM:Gly in axons of septal GAD2+ cells in CA1. Line scans in grayscale across layers (PSAM vs GFP, unpaired *t*-test, P = 0.01). sa, stratum alveus; so, oriens; sp, piramidale; sr, radiatum; slm, lacunosum-moleculare. (**c**) Local chemogenetic inhibition of MS^GABA^ inputs within CA1 at retrieval test (PSAM+PSEM, n = 11; PSAM+VEH, n = 9; GFP+PSEM, n = 6; effect of treatment F_(2, 26)_ = 22.4, P < 0.0001; ANOVA followed by Tukey’s test, PSEM vs VEH, P < 0.0001; GFP vs PSAM, P = 0.0005; GFP vs VEH, P = 0.738). (**d**) Global inhibition of MS^GABA^ cells at retrieval test (vector, n = 8; hM4D, n = 9; effect of treatment F_(2,15)_ = 16.6, P = 0.001; effect of conditioning F_(1,15)_ = 94.8, P < 0.0001; interaction F_(2,15)_ = 16.6, P = 0.001; two-way RM ANOVA followed by Sidak’s test, vector vs hM4D at retrieval, P < 0.0001). (**e**) pErk labeling in CA1 pyramidal cells after vehicle or PSEM infusion during retrieval in mice expressing PSAM as in (a). (**f**) pErk quantification 15 min after retrieval test in the three different treatment groups (effect of treatment F_(5,44)_ = 10.1, P < 0.0001; ANOVA followed by adjusted Tukey’s test for control comparisons, MS^CA1^ cells, P = 0.0006; MS^GABA^ cells, P = 0.0047; MS^GABA^-CA1 terminals, P = 0.0123).

Separate GAD2-cre mice conditionally expressing hM4Di:mCherry in MS^GABA^ cells also showed a deficit in contextual memory upon systemic CNO injection before retrieval test (Fig. 2d and Supplementary Fig.4d). In contrast, tone-evoked memory, which does not rely on hippocampus, or open field behavior were spared by MS^GABA^ inhibition (Supplementary Fig. 4e-h), ruling out indirect effects on locomotion or a deficit in expressing a normally retrieved memory.

Re-activation of the Erk/MAPK signaling cascade is one of the few established molecular correlates of memory expression, particularly in the CA1 region^8,20^. Levels of the active, dually phosphorylated form of Erk1/2 (i.e. pErk) increased following memory retrieval in CA1, but not in other brain regions targeted by MS^GABA^ axons (Supplementary Fig. 5). Notably, the induction of pErk in CA1 pyramidal cells (i.e. WFS1+) was strongly suppressed when MS^CA1^ neurons, MS^GABA^ terminals in CA1 or MS^GABA^ cells were specifically inhibited compared with controls (Fig. 2e,f), demonstrating that the MS^GABA^– CA1 pathway is instrumental for memory-associated MAPK activation.

To elucidate the microcircuit involved, we obtained acute hippocampal slices from GAD-cre mice previously injected in the MS with AAV-DIOChR2:mCherry (Supplementary Fig. 6a,b). Under voltage clamp, light pulses elicited synaptic currents that were blocked by TTX and revived after the addition of 4-AP, suggesting a monosynaptic connection (Fig. 3a). The response persisted in the presence of CNQX/ D-AP5, and it was only blocked after the addition of gabazine, thereby confirming septal GABAergic input. All (13/13) fast spiking (FS) cells, and 83% (34/41) of non-FS cells showed suppressed firing during optogenetic stimulation (Fig. 3b and Supplementary Fig. 6c,d). In contrast, 6% (3/47) of pyramidal cells and none of the giant radiatum cells (0/3) responded to stimulation. We also injected GAD-cre mice in the MS with an anterograde trans-synaptic virus, AAV1-DIOFlpo, followed by injection in CA1 of AAV5-fDIO-YFP for genetic amplification. Somata expressing Flp recombinase and hence Flp-dependent YFP mainly lied in the pyramidal cell layer and expressed interneuron markers in various proportions (Supplementary Fig. 6e-h); a large subset was parvalbumin-rich (PV+). MS^GABA^ inputs may therefore limit the recruitment of inhibition during successful recall. Consistent with this notion, acute suppression of MS^GABA^ gated the activation of FS/PV+ cells in CA1 alongside with memory deficits (Fig. 3c and Supplementary Fig. 7). Because the role of PV+ cells on memory retrieval is unclear, we conditionally expressed the excitatory receptor hM3Dq:mCherry in dorsal CA1 of PV-cre mice (Supplementary Fig. 8a-c), i.e. mainly in basket and bistratified inhibitory cells^21^. Mice froze significantly less in response to CNO during contextual re-exposure compared with vehicle (Fig. 3d); they also showed a strong deficit in pErk1/2 in pyramidal cells that was inversely correlated with PV+ cell activation (Fig. 3e, and Supplementary Fig. 8d-h). Arguing against nonspecific deficits, the same PV-cre mice showed a modest improvement in a Y-maze spatial working memory task when treated with CNO in a permutated design study performed one week before conditioning (Supplementary Fig. 8i). Together, the results point to CA1 PV+ cells as mediators in the effect of the MS^GABA^–CA1 pathway on contextual memory recall.

**Figure 3.**
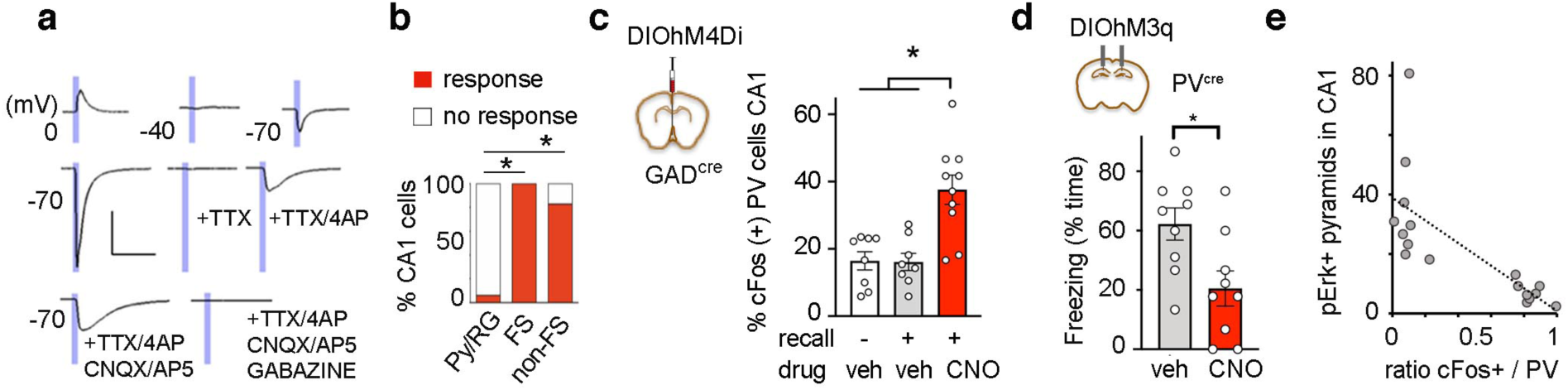
Feed-forward interneurons mediate the effect of MS^GABA^ on memory retrieval. (**a**) Sample response traces recorded at different voltages upon photostimulation of ChR2-mCherry positive septal axons in CA1. The reversal potential of chloride ions shifts at −40 mV with the intracellular solution used (Methods). Sample response traces during consecutive drug applications were recorded at −70mV. Scale, 200 pA, 40 ms. (**b**) Percentage of responding cells upon photostimulation of MS^GABA^ fibers (Fischer’s exact test, excitatory vs FS, P < 10^−10^; excitatory vs non-FS, P < 10^−13^). (**c**) Effect of MS^GABA^ inhibition during retrieval on c-Fos in CA1 PV+ cells (effect of treatment F_(2,25)_ = 12.8, P = 0.0002; ANOVA followed by adjusted Tukey’s test, naïve vs recall in VEH, P = 0.99; recall in VEH vs CNO, P = 0.0007; naïve vs recall in CNO, P = 0.0008). (**d**) Context-evoked freezing during hM3q stimulation of CA1 PV+ cells in PV-cre mice (VEH, n = 9; CNO, n = 9; unpaired *t*-test, t = 2.7, P = 0.015). (**e**) Inverse correlation between pErk+ pyramidal cells and cFos+ PV cells following retrieval test in PV-cre mice (control and hM3q-stimulated; r = - 0.72, t = - 3.72, P = 0.002).

We provide direct evidence that expression of contextual fear behavior in mice is causally related to activation of a septal GABAergic projection to the CA1 hippocampus, restricting local inhibitory control to enable activation of MAPK signaling in memory-holding pyramidal cells^4,6,8,20^ (Supplementary Fig. 9). By encoding the intensity of multisensory cues^15^, septo-hippocampal GABAergic boutons may calibrate behavioral or autonomic responses^12^ upon contextual re-exposure. Further, the enhanced response of MS^CA1^ cells to conditioned stimuli following learning parallels observations in the parabrachial nucleus^22^, and suggests a mechanism underlying the efficiency of memory recall. The ability of the MS^GABA^–CA1 pathway to prevent the induction of PV+ cell activation during retrieval is also analogous to the disinhibition of excitatory outputs in prefrontal cortex that gates fear expression^23^. This role of MS^GABA^–CA1 projections is qualitatively different from the excitatory septal outputs that encode innate valence^24^, but together highlight the capacity of the MS to orchestrate learned and innate emotional responses through anatomically segregated pathways.

## Methods

### Animals

Procedures involving mice were performed in accordance with EU guidelines (2010/63/EU), and had ethical approval from the animal and ethical committee at University of Barcelona, and from the Regierungspräsidium Karlsruhe, Germany (G-254-14). Adult C57BL/6J mice were supplied by Charles River. GAD2-Cre and PV-cre mice of either sex and expressing Cre recombinase in glutamic acid decarboxylase or parvalbumin positive cells were bred in-house on a C57BL6 background, and used at 2-5 months of age. Mice had unrestricted access to water and food and maintained on a 12 h light cycle.

### Viral constructs

Herpes viruses (HSV-YFP, HSV-YFP-IRES-cre, HSV-mCherry-IRES-cre) were supplied by the MIT Viral Gene Transfer Core at titers of 3.5 × 10^8^ IU/mL. CAV2-cre virus was provided by Eric Kremer and amplified in-house to ~10^12^ vg/mL. Adeno-associated vectors AAV8-EF1a-DIO-hM4Di-mCherry, AAV5-EF1a-DIO-hChR2(H134R)-EYFP, AAV5-EF1a-DIO-EYFP, and AAV1-CAG-flex-tdTomato were supplied by the UNC vector core facility at titers of 4-7 × 10^12^ vg/mL. AAV1/2-hSyn-flex-PSAM^L141F^:GlyR-IRES-GFP (addgene plasmid #32479) was produced in-house by calcium phosphate-mediated transfection of 293T cells, purified by sucrose and CsCl gradient centrifugation steps, and resuspended in HEPES buffered saline to ~10^12^ vg/mL. AAV1/2-DIO-hChR2(H134R)-mCherry was obtained from Karl Deisseroth. AAV5-hSyn-DIO-hM3Dq-mCherry was supplied by addgene (#44361) at 7 × 10^12^ vg/mL. AAV1-EF1a-DIO-FLPo (addgene plasmid #87306) was produced by Vigene Biosciences at 2 × 10^13^ vg/mL. AAV5-CAG-dFRT-EYFP was supplied by the viral vector facility at Neuroscience center Zurich at 1 × 10^13^ vg/mL.

### Stereotaxic surgery

Mice were anesthetized with ketamine/xylazine and placed in a small-animal stereotaxic instrument (WPI). Detailed procedures followed a previous protocol^25^. Eyes were lubricated with ophthalmic ointment, fur shaved, skin sterilized and bupicaine locally administered at incision site. A small craniotomy was followed by viral injection or Alexa 488-conjugated cholera toxin-B with a Hamilton syringe (33 gauge) at a rate of 35 nL / min for a total of 300-800 nL. Injection coordinates are given in mm, relative to Bregma and brain surface, for dorsal CA1: −1.9 AP, 1.4 ML, −1.45 DV; for MS: +1 AP, 0 ML, −4 DV; for EC: −4.65 AP, 3.25 ML, DV −2.8. Recovery times before behavior were optimized for each virus: 2 weeks for HSV, 4-5 weeks for AAV-PSAM, 6 weeks for CAV2, 3-4 weeks for other AAVs. For intra-hippocampal drug infusion or septal photostimulation, custom-made cannulae (30 G; Plastics1, Inc.) or fiber-optic cannulae (200 um diameter; Doric Lenses) were implanted after virus injection in the same surgery, and secured with dental acrylic and microscrews.

### Electrophysiology

GAD-cre mice were stereotaxically injected with 300 nl of AAV1/2-DIO-hChR2(H134R)-mCherry via a glass micropipette (tip resistance of 2 to 4 MΩ). The scalp incision was sutured and the mice were constantly monitored for proper recovery. Two weeks post-surgery, mice were deeply anaesthetized with inhaled isoflurane, followed by transcardial perfusion with ~20 ml ice-cold sucrose solution containing (in mM) 212 sucrose, 26 NaHCO3, 1.25 NaH2PO4, 3 KCl, 7 MgCl2, 10 glucose and 0.2 CaCl2, oxygenated with carbogen gas (95% O2/ 5% CO2, pH 7.4). 300 µm sections were cut in ice-cold oxygenated sucrose solution, followed by incubation in oxygenated extracellular solution containing (in mM) 12.5 NaCl, 2.5 NaHCO3, 0.125 NaH2PO4, 0.25 KCl, 2 CaCl2, 1 MgCl2 and 25 glucose. Individual slices were placed in a submerged recording chamber mounted on an upright microscope (Olympus BW-X51) and continuously perfused with oxygenated extracellular solution. Cells in CA1 of the hippocampus and in MS were visualized with DIC optics and epifluorescence was used to detect mCherry fluorescence.

We recorded from acute horizontal slices. The septal injection site was controlled in coronal sections. Recording pipettes were pulled from borosilicate capillaries with the tip resistance of 4-6 MΩ and filled with an intracellular solution containing (in mM): 105 K-gluconate, 30 KCl, 10 Hepes, 10 phosphocreatine, 4 Mg-ATP and 0.3 GTP, pH adjusted to 7.3 with KOH. This intracellular solution causes the reversal potential of the chloride ions passing through the GABA channel to shift to −40mV thereby allowing GABAergic inputs to be observed better. Liquid junction potentials were not corrected for. Target cells in different layers of CA1 were patched and classified based on firing paterns^25,26^. To analyze the postsynaptic responses, cells were voltage-clamped at −70mV and were stimulated with 5ms LED-stimulation at 470nm. Bath-applied drug concentrations were: CNQX (10 µM), D-AP5 (50 µM), gabazine (10 µM), TTX (1 µM) and 4-AP (100 µM). All recordings were made using HEKA PatchMaster EPC 10 amplifier, signals were filtered at 3 kHz, and sampled at 20 kHz. Data were analyzed offline with MatLab and results are presented as mean± SEM.

### Fear conditioning

For context-evoked memory, mice were placed in a standard fear conditioning box (Coulbourn Instruments) wipe-cleaned with 1 % acetic acid, 40 dB white noise, and indirect room light. Following 2 minutes of exploration, they received a single 2 s 0.7 mA foot shock and removed 58 s later. One or two days later, they were placed back into the same context for 3 minutes to assess freezing. Naïve mice were never shocked. Behavior was automatically recorded with Smart software (Panlab) and freezing hand-scored every 5 s by blinded experimenters. For delayed tone conditioning, three 0.3 mA 2 s shocks were co-terminated with 15 s 2 kHz, 80 dB tones delivered in succession at 60-90 s intervals. On day later, mice were placed in a different context (with bedding, round corners, opaque walls cleaned with 10 % ethanol) that evoked no freezing behavior until the same 15 s tone was delivered 4 times at 1 min intervals. Freezing was scored every 2 s during each tone and the following 15 s.

For laser stimulation, mice had been acclimated for 3 days to fiber-optic cables. During photostimulation experiments, light pulse trains (10 ms pulses at 10 Hz for 30 s) were programmed with a waveform generator (Agilent Technologies, model 33500B) that provided input to a blue light DPSS laser (473 nm; LaserGlow). Light power exiting the fiber-optic cable was ~15 mW under continuous mode. Light trains were delivered 3 times at 2 min intervals during the retrieval test.

### Open field

Mice were placed in an arena (40 x 40 x 40 cm), and allowed to explore for 5 min while behavior was recorded with Smart software. Center zone (25 % area) was predefined. Move transitions were defined as movement events following a > 2 s resting time (i.e. < 0.5 cm / s).

### Y-maze

A discrete alternation version was used. Every trial consisted of two runs, and trials were repeated 6 times at 20 min intervals. In the first run of a given trial, the mouse was placed in the holding compartment that was connected through a sliding door to the start arm of a Y-maze (10 x 24 x 8 cm arms). The mouse was allowed to choose one of the two distal target arms, after which it was enclosed for 30 s. It was then returned to the holding chamber, and 5 s later allowed to run the maze again to record the target arm chosen (i.e. the same arm or the new, alternative arm).

### Pharmacological injections

CNO (3 mg/kg, intraperitoneal injection 30 min before test; Enzo Biosciences) was prepared in sterile 0.9% saline and 0.5 % DMSO; PSEM-308 was prepared at 100X concentration in nuclease-free water, and diluted to 100 uM in aCSF the day of use. Intrahippocampal infusion of PSEM or vehicle was performed 10 min before the test, with cannulae connected to Hamilton syringes through polyethylene tubing and mounted in an infusion pump. A subset of mice (n = 8) was co-injected with 0.5 % Alexa555-conjugated dextranamine to verify drug diffusion. The mouse was unrestrained and conscious during infusion (0.5 µl / side, 0.2-µl/min), which was allowed to diffuse for 2 additional minutes before re-inserting dummy cannulae, and returning to the home cage.

### Tissue processing and staining

Mice were deeply anesthetized and transcardially perfused with ice-cold saline, followed by 4% formaldehyde in phosphate buffer (PB) 0.1 M. Na_3_VO_4_ (1mM) and NaF (50 mM) were added to saline and formaldehyde solutions when p-Erk1/2 immunostaining was planned. Brains were post-fixed overnight (ON) at 4 ºC and transferred to 30% sucrose solution in PB until they sunk. Brains were frozen in dry ice and sliced at 25 um thickness in a cryostat. Serial sections were kept in cryoprotector solution at −20ºC until use. In most instances, virally expressed fluorophores were visualized without immuno-amplification. Sections were permeabilized with 0.3% TritonX-100 or 0.1% Tween-20 in buffer solution, and incubated free-floating in antibodies with 10% serum and 2 % BSA ON at 4ºC. Primary antibodies were mouse anti-GAD67 (1:500) and goat anti-ChAT (1:250) from Millipore; guinea pig anti-VGAT (1:250), and guinea pig anti-RFP (1:500) from Synaptic Systems; rabbit anti-pErk1/2 #9101 from Cell Signaling at 1:300 (for bright field) or 1:10,000 (for tyramide amplification); rabbit anti-parvalbumin, anti-calbindin, anti-calretinin (all at 1:5,000 from Swant); rabbit anti-somatostatin, anti-cholecystokinin-8, anti-vasointestinal peptide (all at 1:2,000 from Immunostar); goat or rabbit anti-cFos (# sc-52 at 1:1,000 from Santa Cruz); rabbit anti-WFS1 (1:500) from Proteintech. The next day, the tissue was washed and incubated for 2 h in species-specific secondary antibodies conjugated to Alexa 488/555/647 (1:500, Invitrogen) or biotin (1:250, Vectorlabs). Tissue with biotinylated antibodies was further incubated with HRP-conjugated streptavidin (1:250, Perkin Elmer) and visualized either with DAB or FITC-conjugated tyramide (Perkin Elmer) following manual instructions.

### Imaging and analysis

Surgery cases included in the formal analysis were individually verified for successful viral expression, drug infusion and anatomical location. Images were acquired using a Leica SP5 confocal microscope, or a Leica AF6000 microscope for low magnification or bright field. For comparisons of behavioral effects, laser power, spectral range, pinhole aperture, detector gain, and offset were initially set to obtain pixel densities within a linear range, and were then kept constant between experimental groups or brain regions. To estimate the fraction of cells co-labeled for cFos, pErk or a type-specific marker, Z-series stacks of 1024 x 1024 pixel images (average of 4 scans per plane) were typically obtained from at least 2-3 consecutive sections per mouse (125 ums apart) with a 40X objective, thus obtaining a mean value per subject. Co-labeling in somata was assessed near the midplane of individual cells based on DAPI counterstaining. To estimate the fractional CA1 input among afferent regions ~1,000-2,000 YFP+ cells / series of sections was recorded per mouse. The neurochemical profile of MS^CA1^ cells was estimated from 345 YFP+ cells and 5 mice. The fractions of Fos+ MS^CA1^ cells were estimated from 668 YFP+ cells and 19 mice; the neurochemical profile of hippocampal interneurons postsynaptic to MS was estimated from 254 GFP+ cells and 3 mice; the fractions of Fos+/PV+ cells after MS^GABA^ inhibition were estimated from 536 cells and 25 mice; 3,364 pErk+ cells in dorsal CA1 were analyzed from 44 mice with MS perturbations.

For quantification of axonal projections, z-stacks were collapsed, background-substracted, thresholded, minimally filtered, and axons automatically selected in ImageJ; the created selection was then overlapped over the original image using the ROI function, and the integrated intensity within was measured in a 100 x 100 um square in each of the examined regions. In all cases, image acquisition was done blind to treatment group, and sampled regions were determined based on precise anatomical identification, tissue preservation and/or presence of the cells to be interrogated; the channel reporting the dependent measure was then visualized and recorded.

### Statistics

Prism 8.0 (GraphPad) was used for statistical analysis. Two-tailed, unpaired *t*-tests were used when comparing two data sets. To analyze multiple groups or regions, one-way ANOVA, two-way ANOVA with post hoc Tukey’s tests, or repeated measures ANOVA with post hoc Sidak’s tests were used as appropriate. Sample sizes (n > 6) for 90% power and alpha = 0.05 considered the mean ± SD of freezing behavior previously observed in WT conditioned mice in our laboratory (65 ± 16 %, n = 27). A behaviorally relevant magnitude of difference was set to > 30% based on previously published pharmacological effects on retrieval^8^. Behavioral effects were validated through complementary approaches, and further correlated with corresponding molecular effects, arguing against potential type I errors. Data sets were large enough to confirm with Shapiro-Wilk test and QQ plots that they followed a normal distribution. Error bars represent means ± standard error. All *n* values represent individual mice.

## Supporting information

supplementary data_SansDublanc

## Contributions

A.S., A.R., S.D. and C.S. performed and analyzed experiments. H.M. provided reagents and supervised electrophysiology experiments. M.P. and J.R. provided reagents and contributed to the experimental design. C.S. conceived and coordinated the project, and wrote the manuscript. All authors contributed interpretation of the data and commented on the manuscript.

## Acknowledgements

We thank Scott Sternson, Li Zhang, Brian Roth, and Karl Deisseroth for depositing plasmids to Addgene. Funded by grants SAF2012-40102, SAF2015-69767P (MINECO/FEDER), and Marie Curie CIG-631035 to C.S.

