## supplementary data_SansDublanc for "Septal GABAergic Inputs to CA1 Govern Contextual Memory Retrieval"

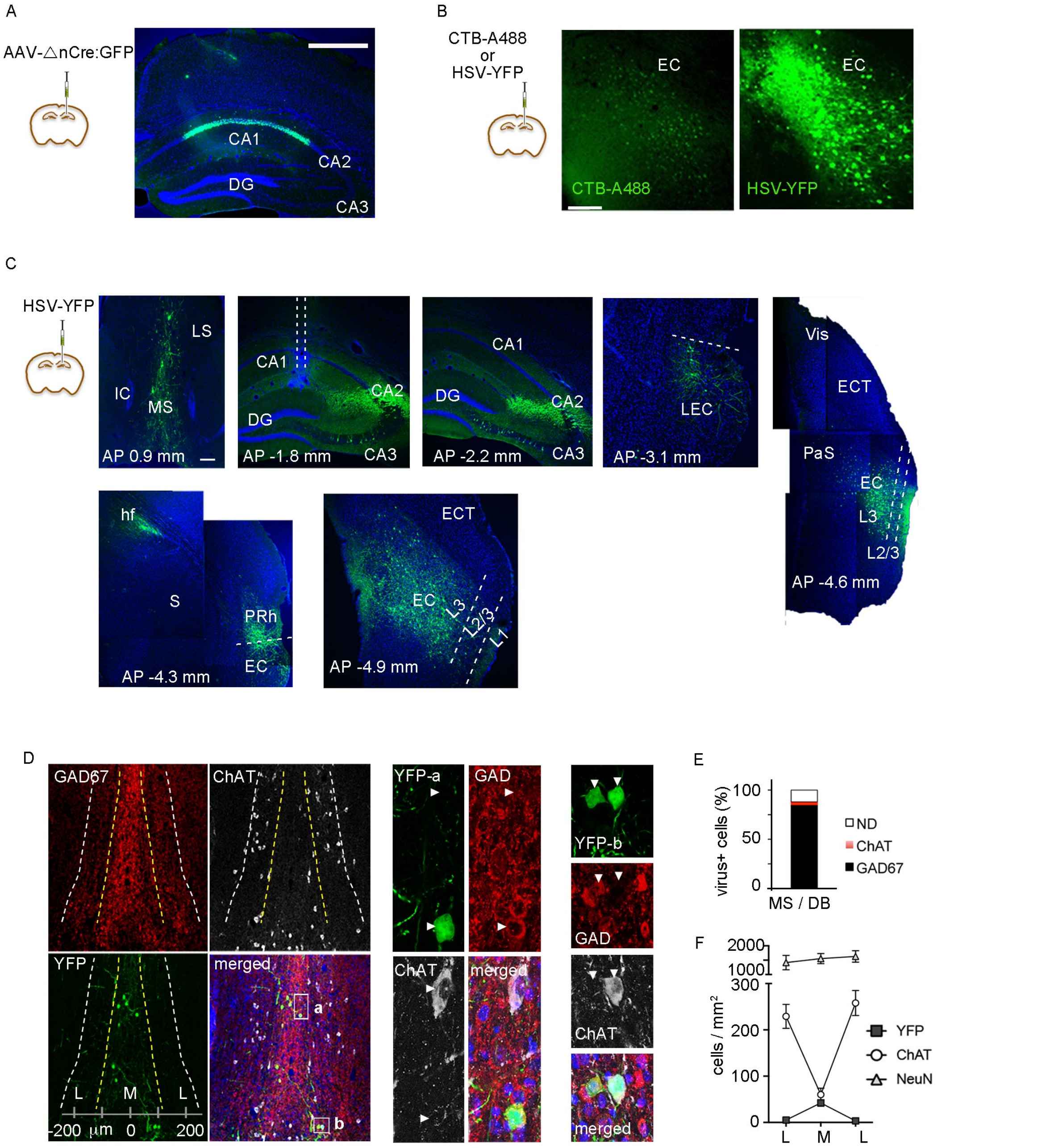


### Supplementary Figure 1

CA1-projecting cells in the brain

(a) Stereotaxic injection of inactive nuclear Cre:GFP (AAV-△nCre:GFP) showing transduction confined to CA1. Scale bar, 0.5 mm.

(b) Comparison of retrograde labeling in entorhinal cortex (EC) after injection in CA1 of Alexa488-conjugated cholera toxin B or HSV-YFP, the latter being an order of magnitude stronger without amplification. Scale bar, 100 um.

(c) Examples of retrogradely labeled regions in the brain after HSV-YFP injection in CA1. IC, islands of Calleja; LS, lateral septum; MS, medial septum; DG, dentate gyrus; LEC, lateral entorhinal cortex; Vis, visual cortex; ECT, ectorhinal cortex; PaS, parasubiculum; hf, hippocampal fisure; S, subiculum; PRh, perirhinal cortex; L, layer. Scale bar, 100 um.

(d) Confocal images of the medial septum co-labeled for GAD-67 (red) and ChAT (silver) in a mouse injected with HSV-YFP in CA1. Small rectangles in the merged image correspond to magnified images on the right.

(e) Quantification of CA1-projecting septal cells co-labeled for GAD or ChAT; ND, non-determined.

(f) Density estimates of cells immunoreactive for ChAT, NeuN or retrograde YFP in central (M) and lateral (L) divisions of MS from midline.

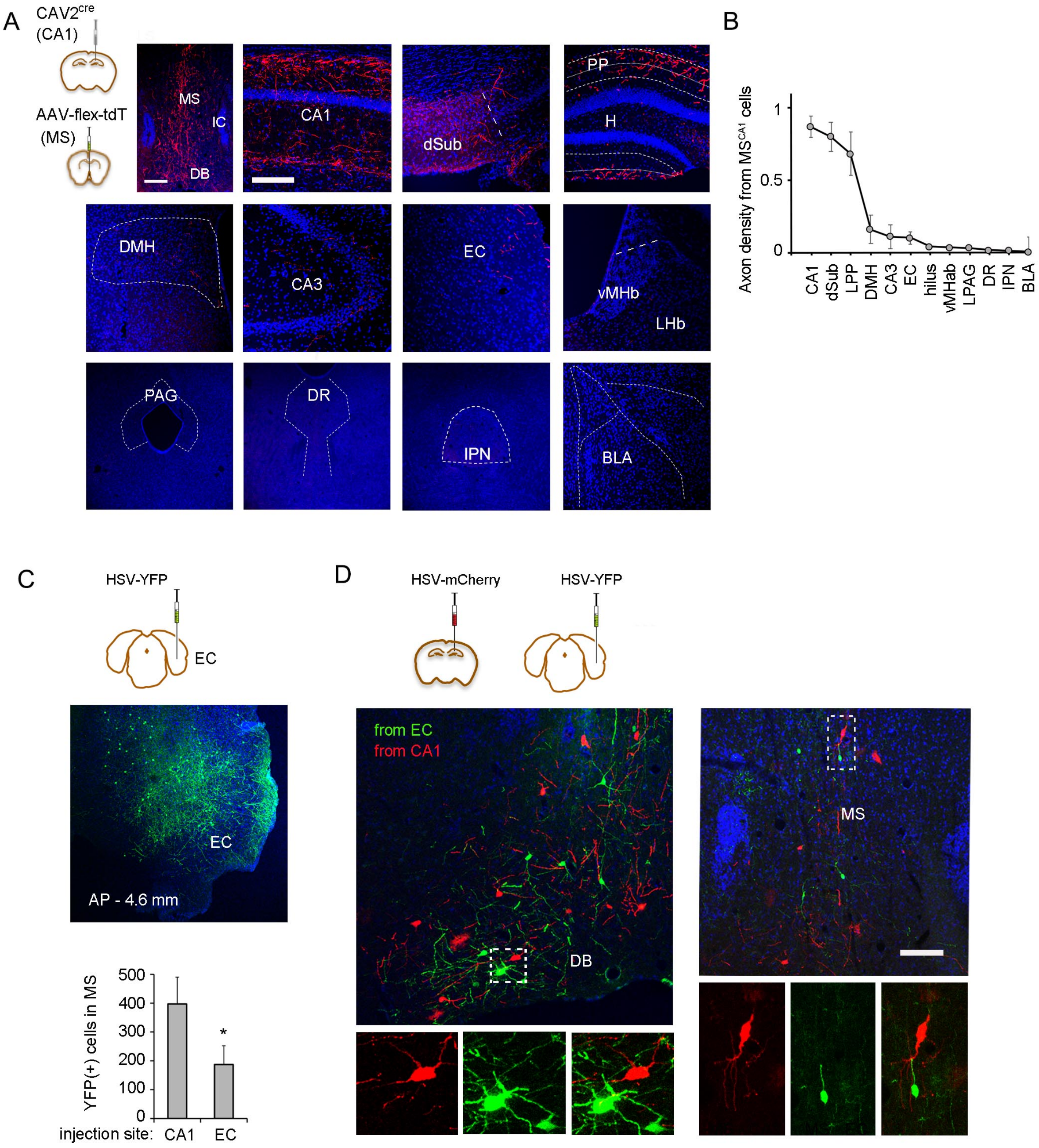


### Supplementary figure 2

Collateralization of CA1-projecting septal cells.

(a) Confocal images showing the axonal distribution of CA1-projecting septal cells conditionally labeled with tdTomato in various brain regions. DB, diagonal band; dSub, dorsal subiculum; PP, perforant path; H, hilus; DMH, dorsomedial hypothalamus; vMHb, ventral division of medial habenula; LHb, lateral habenula; PAG, periaqueductal gray; DR, dorsal raphe; IPN, interpeduncular nucleus; BLA, basolateral amygdala. Scale bar for MS, 100 um. Scale bar for target regions, 100 um.

(b) Quantification of axonal densities normalized to that in CA1. Axonal density across regions (n = 3; F(1.8, 3.6)= 15.5, P = 0.016).

(c) Injection of HSV-YFP into the EC, and quantification of retrogradely labeled cells in the MS after injection in EC vs CA1 (n= 6 / group; *t*-test, t = 4.6; P = 0.001).

(d) Co-injection of HSV encoding mCherry (in CA1) or YFP (in EC) labels distinct cell populations in the MS complex. Scal bar, 100 um.

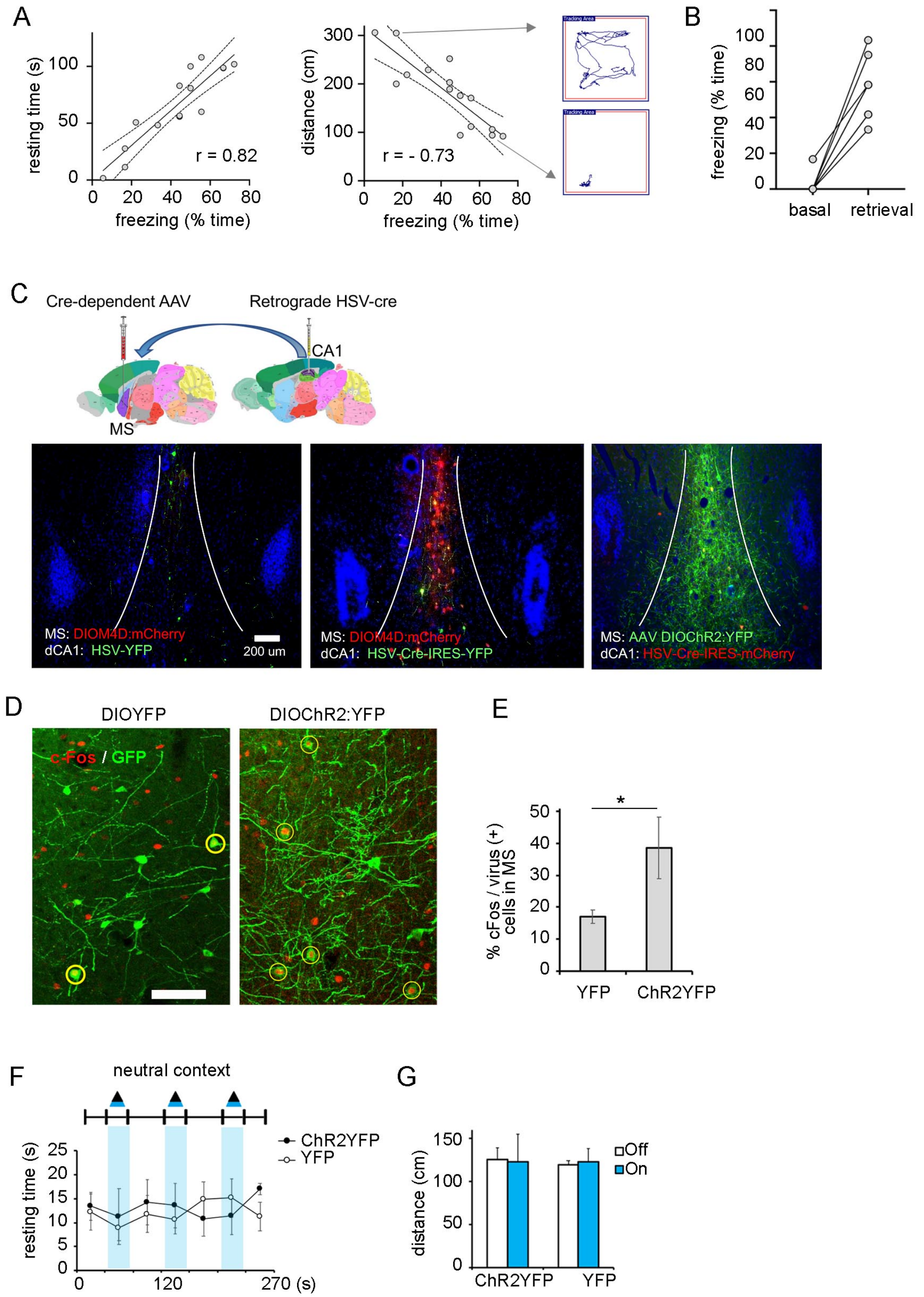


### Supplementary Figure 3

Functional targeting of the MS-to-CA1 pathway

(a) Correlation between freezing scores and automatically recorded (center body point) resting time or ambulatory distance. Example track lengths from individual mice with high or low freezing shown on the right. (b) Freezing behavior before conditioning and upon re-exposure to context aftert conditioning. (c) Scheme of viral injections used to express recombinant actuators in $MS^{CA1}$ cells, and examples below confirming the cre-dependent, projection-specific expression of hM4D or ChR2.

(d) Images of MS following retrieval paired with photostimulation YFP or ChR2YFP-expressing cells. (e) Quantification of cFos (two-tailed t-test, $t = 3.45$, $P = 0.007$). (f-g) Photostimulation of $MS^{CA1}$ spares resting time and ambulatory distance in an open field.

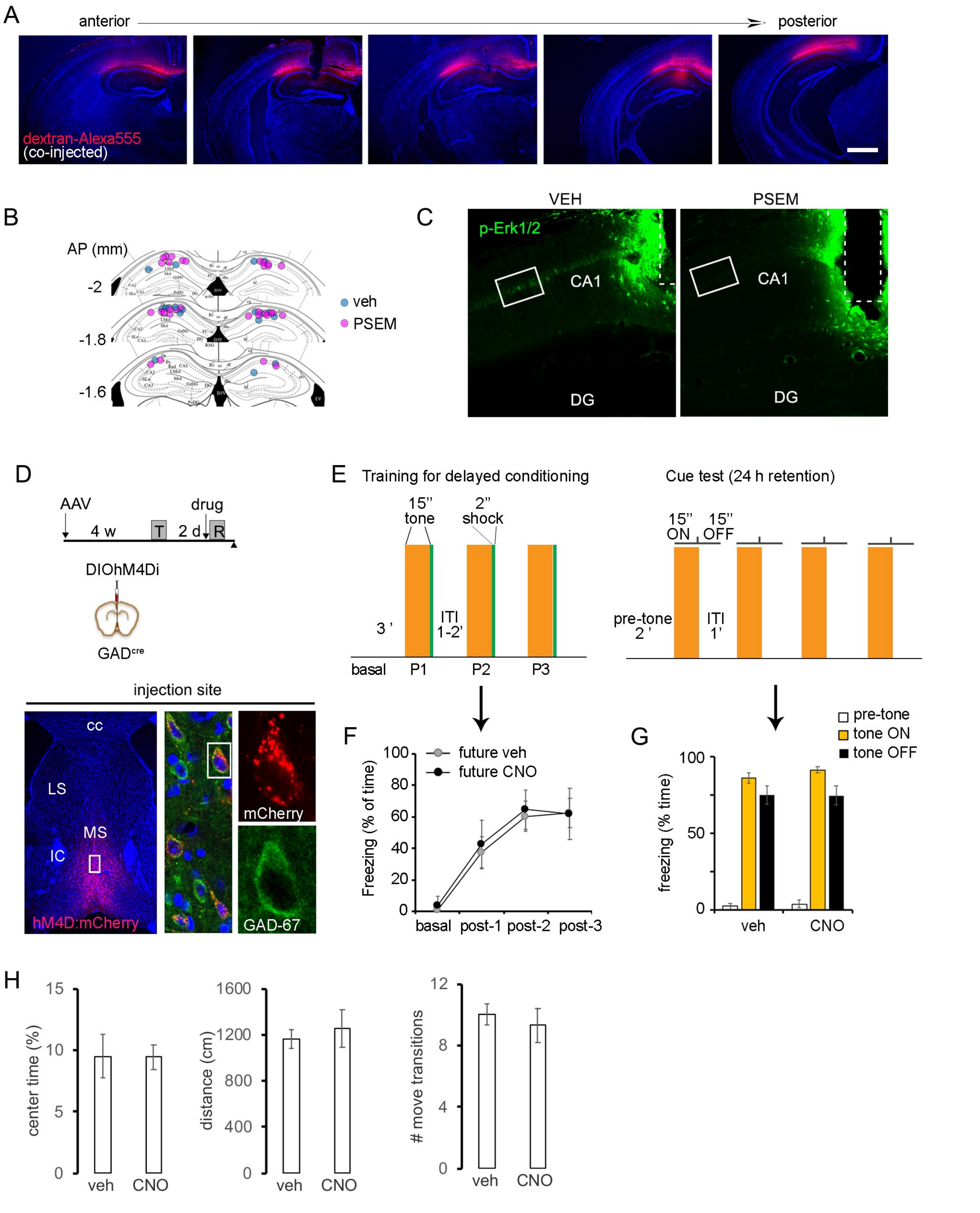


**Supplementary figure 4**

Local or global targeting of septal GABAergic cells.

(a) Example of a mouse injected with PSEM together with fluorescent dextranamine along the antero-posterior axis of the hippocampus.

(b) Scheme of bilateral cannulations in mice infused with PSEM or vehicle.

(c) Low-magnification image of p-Erk fluorescence. Rectangles correspond to locations analyzed for pErk quantification (> 200 um away from cannula track).

(d) Experimental design used in experiments targeting the general septal GABAergic population, and representative example below.

(e) Auditory delayed conditioning paradigm for training and memory test.

(f) Freezing curves during conditioning.

(g) Mean freezing values during the cue test in vehicle (n = 8) and CNO-treated (n = 9) mice (t-test veh vs CNO tone on, t= 1.5, P = 0.17).

(h) Mean values in the open field for vehicle and CNO-treated mice (n= 6 / group; center time, P = 0.47; distance, P = 0.92; transitions, P = 0.18).

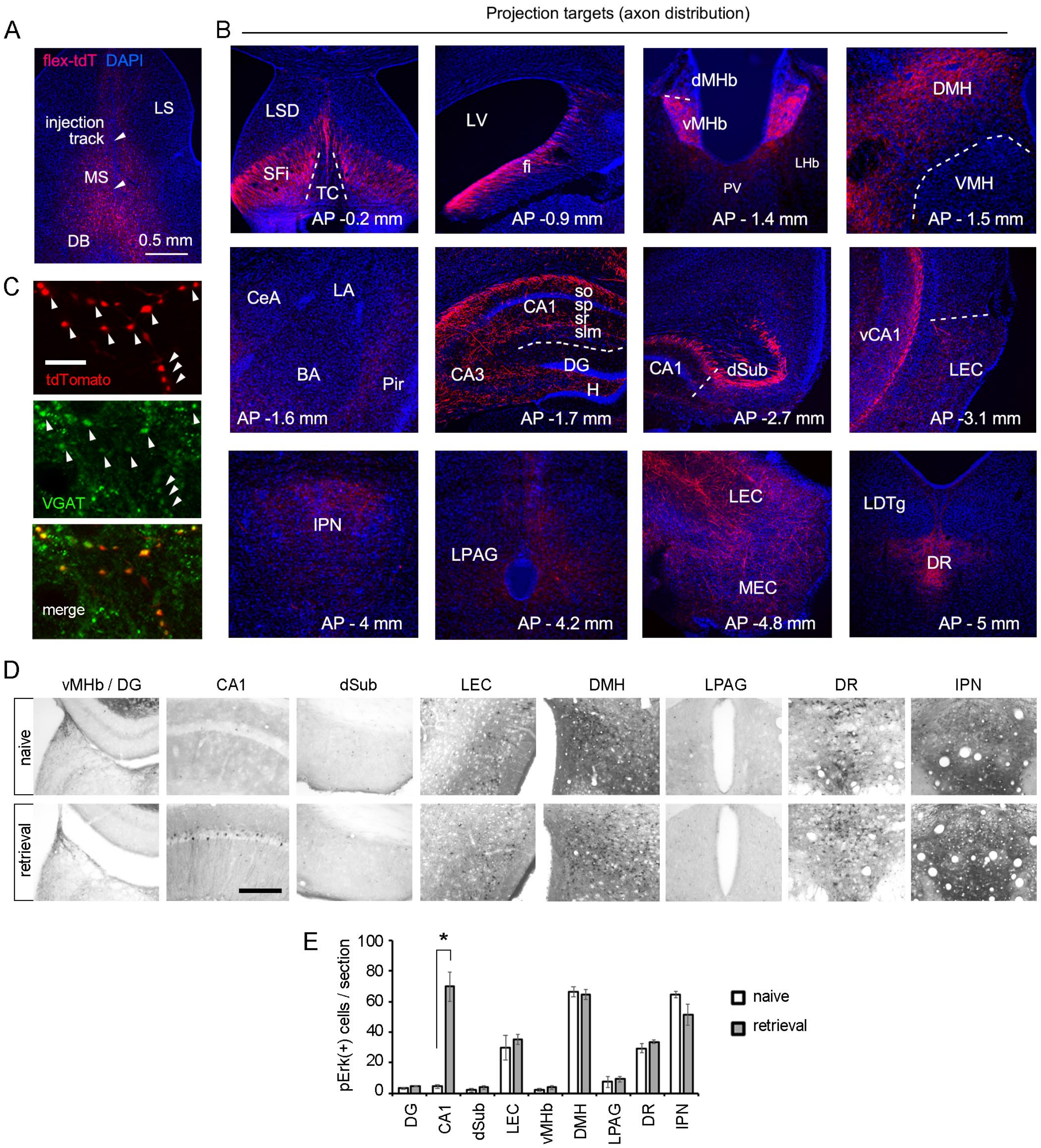


**Supplementary figure 5**

Distribution of input fibers and phospho-Erk1/2 in regions targeted by GABAergic septal cells.

(a) Example of the MS from a GADcre mouse at the site of AAV-flex-tdTomato injection.

(b) Septal GABAergic axons expressing tdTomato in multiple brain regions. TC, triangular septal nucleus; SFi, septofimbrial nucleus; LSD, dorsal part of lateral septum; fi, fimbria; PV, paraventricular thalamus; VMH, ventromedial hypothamalus; CeA, central amygdala; BA, basal amygdala; LA, lateral amygdala; Pir, piriform cortex; vCA1, ventral CA1; MEC, medial entorhinal cortex; LDTg, laterodorsal tegmental nucleus.

(c) Septal GABAergic boutons in CA1 co-labeled for the vesicular GABA transporter VGAT.

(d) DAB-developed sections immunoreacted for p-Erk1/2 in WT naive mice (n = 6) or after retrieval test (n = 11). Scale bar, 200 um.

(e) Quantification of pErk in brain regions examined (*t*-test naive vs retrieval in CA1, t= 6.63, P= 0.00005; P > 0.2- 0.74 in other brain regions).

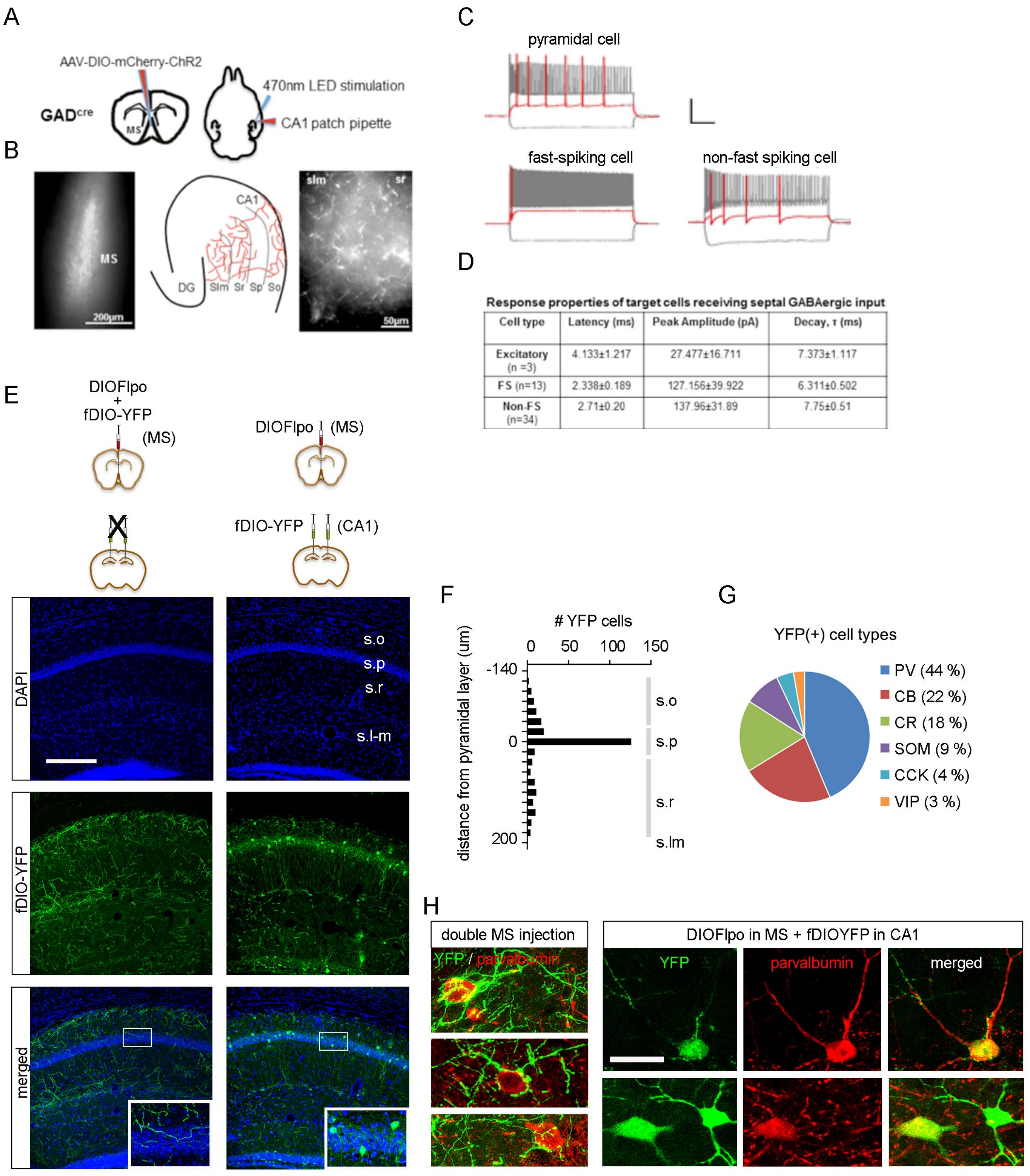


**Response properties of target cells receiving septal GABAergic input**

| Cell type | Latency (ms) | Peak Amplitude (pA) | Decay, τ (ms) |
|---|---|---|---|
| Excitatory (n =3) | 4.133±1.217 | 27.477±16.711 | 7.373±1.117 |
| FS (n=13) | 2.338±0.189 | 127.156±39.922 | 6.311±0.502 |
| Non-FS (n=34) | 2.71±0.20 | 137.96±31.89 | 7.75±0.51 |

### Supplementary figure 6

Hippocampal cells postsynaptic to septal inputs

(a) Schematic diagram of virus injection site in the MS (left); horizontal brain slice showing the placement of the patch pipette and the LED stimulation rod (right). (b) Differential interference contrast image of MS on a coronal brain slice (left). Schematic diagram of the mCherry positive fibers distributed throughout CA1 layers (middle). Sample image of mCherry fluorescence in slm and sr layers within CA1 (right).

(c) Sample firing patterns of different excitatory and inhibitory cells in CA1. (d) Electrophysiological properties of the responses, latency (in ms), peak amplitude (in pA) and decay, $\tau$ (in ms) with mean ± SEM. The response properties of the excitatory cells are significantly different from that of the inhibitory cells. (e) Co-injection of conditional Flpo and Flp-dependent YFP in the MS of GAD-cre mice gives rise to fibers in hippocampus (left), whereas relocating fDIO-YFP injection to CA1 reveals cell somata. Scale bar, 200 um. (f) Distribution of YFP+ cells across CA1 layers.

(g) Proportion of YFP+ cells co-labeled for different interneuron markers in CA1. PV, parvalbumin; CB, calbindin; CR, calretinin; SOM, somatostatin; CCK, cholecystokinin; VIP, vasointestinal peptide. (h) Examples of PV+ cells in CA1 contacted by YFP+ $MS^{GABA}$ axons after double injection of DIOFlpo and fDIOYFP in MS of GAD-cre (left), and putative hippocampal PV+ cell somata postsynaptic to MS co-labeled for Flp-dependent YFP (right). Scale bar, 20 um.

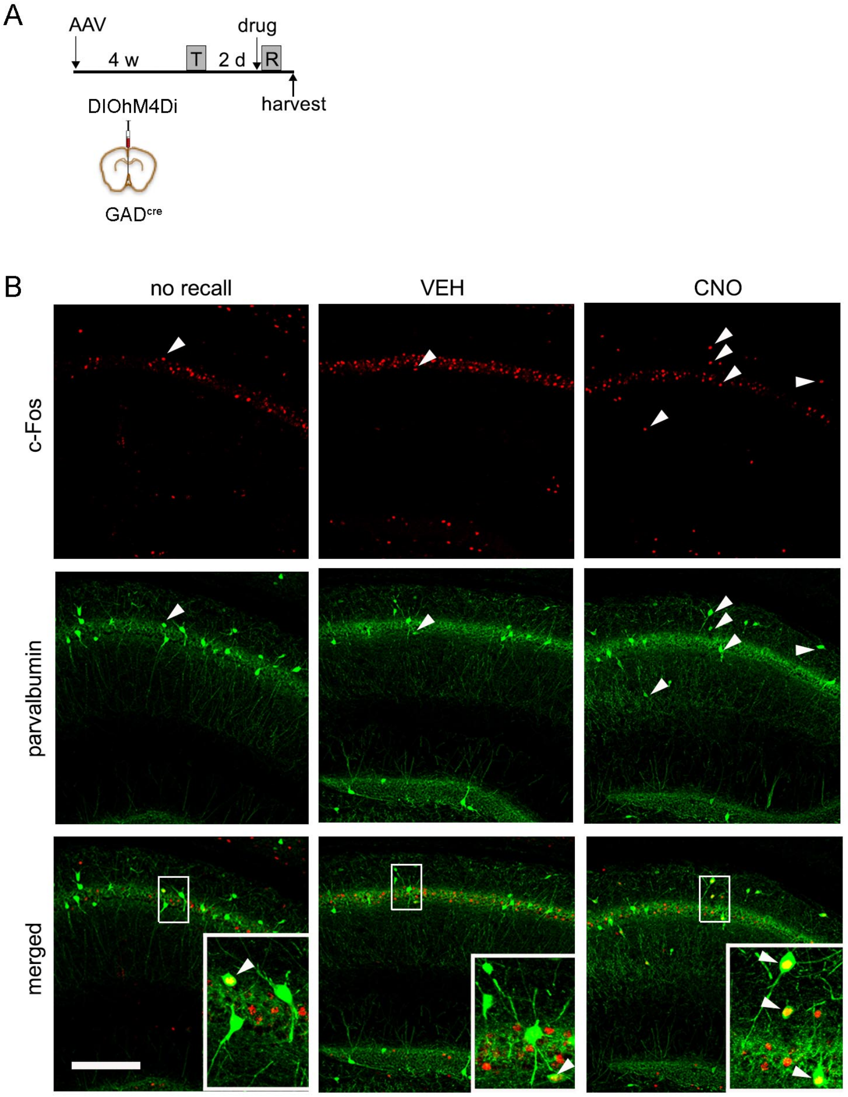


**Supplementary figure 7**

Chemogenetic inhibition of septal GABAergic cells activates parvalbumin interneurons in CA1.

(a) Experimental design.

(b) Representative confocal images of sections double stained for cFos and parvalbumin. Scale bar, 200 um.

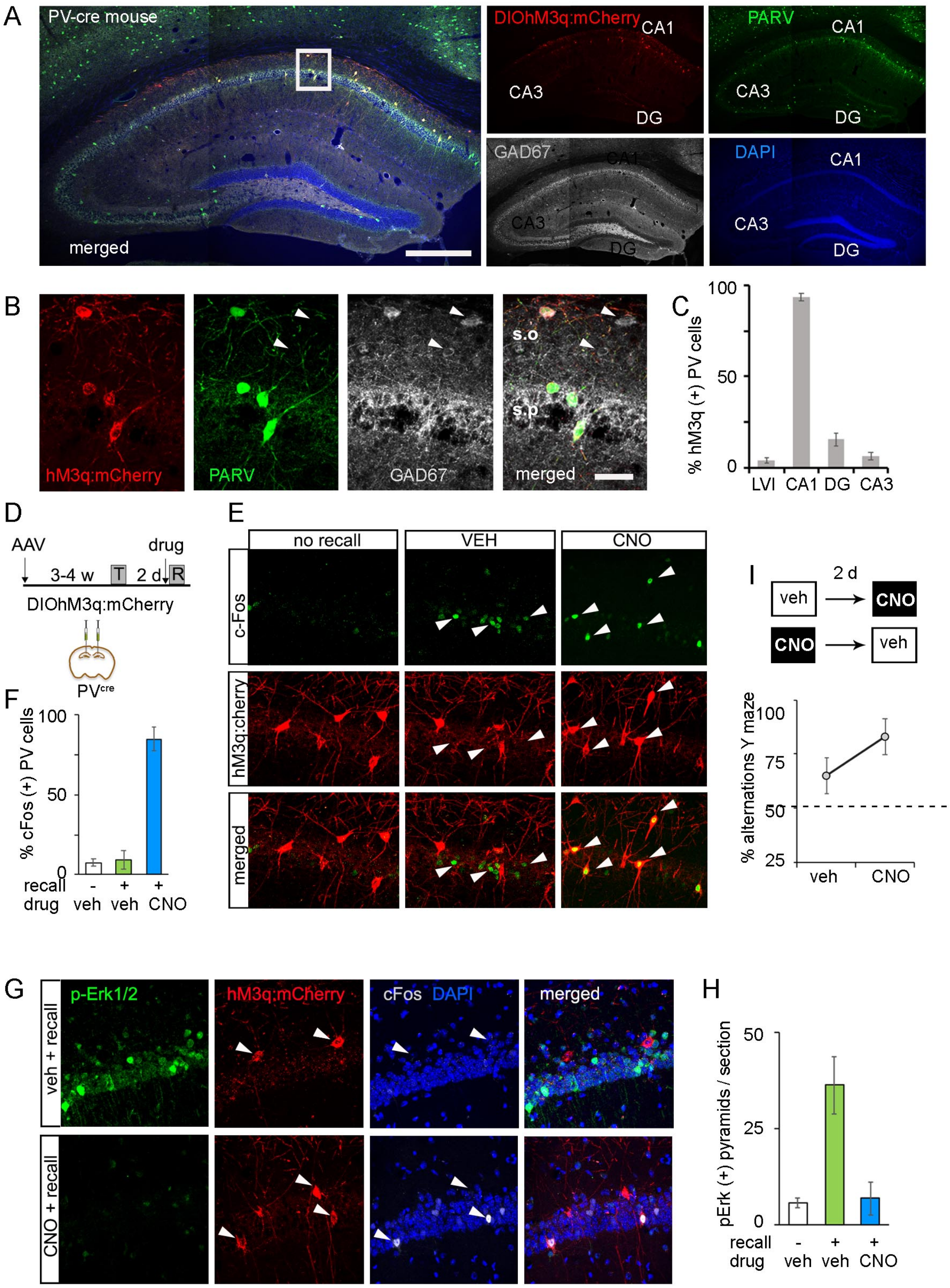


**Suppementary Figure 8**

Conditional expression of hM3q:mCherry in CA1 of PV-cre mice

(a-b) Triple labeling for endogenous parvalbumin, hM3q:mCherry and GAD67 at low (a) and high (b) magnification. Arrowheads point to interneurons negative both for parvalbumin and hM3q. (c) Summary graph showing the fraction of PV cells transduced with hM3q in CA1 and adjacent subfields ($F(3, 38) = 356$, $P < 0.0001$, CA1 vs other subfields, $P < 0.0001$).

(d-f) Experimental design to assess the effect of CA1 PV+ activation on memory recall (d) and pictorial evidence (e) and quantification (f) of corresponding changes in Fos immunoreactivity ($F(2,23) = 723$, $P < 0.0001$; CNO vs VEH groups, $P < 0.0001$).

(i) Summary data of the effect of hM3q activation of CA1 PV+ cells in the Y-maze test. Mice received veh or CNO before the task, and 2 days later they received the alternative treatment before repeating the task (CNO vs VEH, $P = 0.039$, Wilcoxon signed rank test).

(g) Pictorial evidence for the inverse correlation between pErk and parvalbumin activation in CA1. (h) Summary data of pErk+ cells in CA1 of naive, retrieved and retrieved while PV+ cells were hM3q-activated ($F(2,23)=6.1$, $P= 0.012$; recall veh vs CNO, $P= 0.007$)

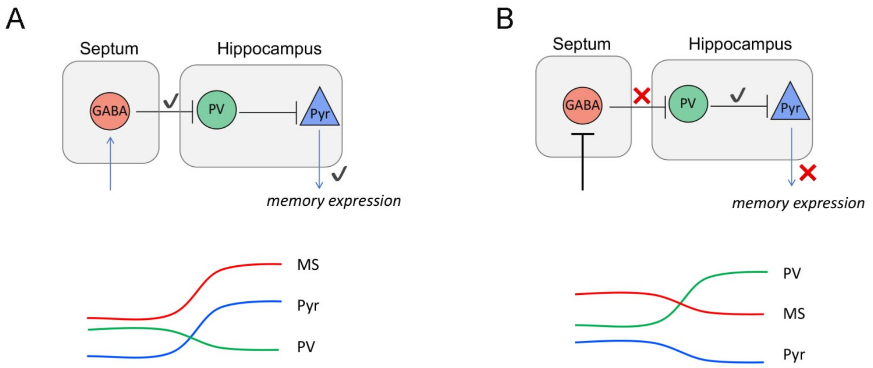


### Supplementary figure 9

Model of septal regulation of contextual memory retrieval

(a) Under physiologcical conditions, activation of septal GABAergic inputs by conditioned stimuli prevents the recruitment of feed-forward interneurons (e.g. PV+ cells), which receive excitatory inputs from Schaffer colaterals [27], thus facilitating signaling events in CA1 pyramidal neurons, such as MAPK activation, critical for memory expression. Reducing disynaptic inhibition via PV+ cells greatly increases the excitatory to inhibitory ratio in CA1 [28]. According to this model, experimental inhibition of CA1 PV+ cells should have little effect on memory recall [29] due to redundancy with MS action.
(b) Experimental blockade of septal GABAergic inputs reveals the permissive role of feed-forward interneurons in retrieval, as PV+ cell disinhibition prevents activation of CA1 pyramidal neurons and memory recall.
